# Circuit Mining in Transcriptomics Data

**DOI:** 10.1101/2025.04.09.647750

**Authors:** Tobias Peherstorfer, Sophia Ulonska, Bianca Burger, Simone Lucato, Bader Al-Hamdan, Marvin Kleinlehner, Till F. M. Andlauer, Katja Bühler

## Abstract

A central goal in neuropharmacological research is to alter brain function by targeting genes whose expression is specific to the corresponding brain circuit. Identifying such genes in large spatially resolved transcriptomics data requires the expertise of bioinformaticians for handling data complexity and to perform statistical tests. This time-consuming process is often decoupled from the routine workflow of neuroscientists, inhibiting fast target discovery. Here we present a visual analytics approach to mining expression data in the context of meso-scale brain circuits for potential target genes tailored to domain experts with limited technical background. We support several workflows for interactive definition and refinement of circuits in the human or mouse brain, and combine spatial indexing with an alternative formulation of sample variance to enable differential gene expression analysis in arbitrary brain circuits at runtime. A user study highlights the usefulness, benefits, and future potential of our work.

---

Neural circuits mediate the brain’s multifaceted functional capabilities at a supracellular level. These circuits consist of interconnected components that collectively serve a specific functional purpose [1], e.g., reward processing. Circuits are often identified from various neuroimaging modalities, e.g. using functional magnetic resonance imaging (fMRI) [2] or cFos stainings [3]. However, circuit function is a complex biochemical process which can be best described and understood by considering multiple *omics* data types at once.

The modification of circuit function using drugs, e.g. to study and treat psychiatric disease, is a main goal in neuropharmacological research. Drug discovery typically begins with drug target identification, where scientists investigate biological entities, such as RNA or proteins, for their suitability as drug targets [4]. In recent years, analyses of spatially resolved transcriptomics data, which quantifies the so-called expression of genes in tissue, have become a key resource in target identification.

A common approach is to mine large bodies of such data for genes that are specific to a circuit of interest, most commonly using so-called differential gene expression analysis (DGEA). Multiple statistical frameworks of varying complexity exist for performing DGEA (e.g., the Wilcoxon signed-rank test or DESeq2 [5]), each contrasting one group of samples to another. Depending on the chosen framework and sample size, computation times range from seconds to several hours [6]. The result of DGEA is a list of genes ranked by the fold change enrichment (ratio of expression between the groups), which can serve as the starting point for searching for molecular targets.

Setting up DGEA pipelines requires bioinformatics knowledge, significant computational resources and time. This is due to the scale and complexity of spatially resolved transcriptomics data (tens of thousands of sample locations with, in commonly used datasets, approximately 20,000 protein-coding genes each). Additionally, multimodal analyses, such as a region of interest (ROI) definition through brain activation data, require the joint integration of transcriptomics data and neuroimaging data in reference brain atlases, which is highly nontrivial. Therefore, neuroscientists or pharmacologists typically cannot easily screen for specific genes in a circuit of interest on their own. A small number of publicly available implementations of such pipelines, such as the Allen Brain web portal [7], exist, but are often tailored towards single datasets and specific types of DGEA analysis, making further extensions challenging.

## Contribution

We present a visual analytics approach for interactive statistical mining of transcriptomics data in arbitrary circuits of the human or mouse brain at the meso-scale (smaller than a brain region, larger than neurons). To the best of our knowledge, this is the first solution enabling neuroscientists without technical background to interactively perform DGEA between arbitrary reference and comparison regions based on spatially resolved transcriptomics data. In particular, we present the following contributions:

- A novel, fully interactive visual workflow for definition and refinement of circuits in the human or mouse brain, allowing the derivation of circuits from brain activation signals and manual selections.
- A novel combination of spatial indexing with an alternative formulation of sample variance, enabling DGEA between arbitrary spatial groups of samples at runtime.
- A visual workflow for performing DGEA using arbitrary reference and comparison groups and interactively filtering query results for further investigation.
- The embedding of our approach into BrainTrawler [8] and its extensive transcriptomic database as well as a small, but novel data resource comprising several pre-defined brain circuits, all made publicly accessible under https://braintrawler-circuit.vrvis.at/

With this work, we do not aim to replace rigorous statistical frameworks for DGEA, but try to provide an efficient way to generate and explore first hypotheses that should be followed up by a more sophisticated statistical analysis. The implementation of DGEA presented in this publication is focused on transcriptomics data, but can be used as-is for multiple other spatial *omics* data types, such as proteomics or ATAC sequencing.

## STATE OF THE ART

DGEA is a standard method in biology with applications in all fields including neuroscience and cancer research [9]. Multiple web tools for performing DGEA on cancer-related data exist. For example, GEPIA2 [10] can be used to query for differentially expressed genes between tumor and normal tissue for multiple cancer-specific datasets. There, users can specify threshold values for specificity and statistical significance before querying and can select between different testing methods. Queries return a table view of the resulting gene list and a visualization of the distribution of result genes on chromosomes. While such applications provide sophisticated statistical testing methods, spatial context is omitted completely. The Allen Brain web portal [7] is an established neuroscience tool that allows spatial comparisons of transcriptomics data. There, users can perform DGEA based on t-tests in one human dataset, with predefined brain regions as the reference and comparison groups. Results are shown as a list and can be sorted by *p*-value and fold change. As an alternative, our previous web tool BrainTrawler [8], [11] allows users to query for highly expressed genes in arbitrary regions and in a multitude of transcriptomics datasets. However, this tool does not use statistical testing in queries and instead only sorts resulting gene lists by the fold-change of expression between the query region and the entire brain. To our knowledge, DGEA using arbitrary user-defined brain circuits is currently not possible with any software. Therefore, we propose an approach to perform interactive DGEA on multi-modal transcriptomics data in arbitrary brain circuits.

## DATA

Before presenting our framework, we briefly summarize which data is being used in our approach and list important characteristics of these data.

### Reference Atlases

Such an atlas consists of a parcellated template brain located in a 3D coordinate space. The parcellation defines how the brain can be split into distinct brain regions, is often organized hierarchically and can be based on cytoarchitecture, functional connectivity, or other modalities. For the human brain, we use the ICBM 152 MNI space template at a resolution of 1mm combined with the Allen Human Reference Atlas parcellation. The Allen Mouse Brain Coordinate Framework at a resolution of 0.1mm is used as a complete reference atlas for the mouse brain. Both reference atlases are established and well-known in the user community. All spatial data is resampled to fit the resolution of the corresponding reference atlas.

### Meso-scale Brain Circuits

A meso-scale brain circuit (smaller than a brain region, larger than neurons) can be defined as a collection of interconnected regions in the brain contributing to a specific function. In this paper, we understand these components as spatially disjoint volumes in 3D brain space and refer to them as nodes. Therefore, we conceptualize a circuit *C* as a set of *n* nodes *c, C* := *{c*_*i*_ *}*_*i*=1..*n*_. Each node *c*_*i*_ is defined by a unique name, a fixed color and a binary mask marking its spatial location and extent. To enable storage and access to predefined brain circuits, we extended the database scheme of BrainTrawler with a tailored data type. Six circuits were derived from Beam *et al*. [12] with the help of domain experts, each of which corresponds to one Research Domain Criteria (RDoC) domain. These circuits were included in the publicly accessible BrainTrawler version.

### Brain Activation Data

Volumetric brain activation data shows which locations in the brain are active for a given task or treatment. Such data can be acquired from, e.g., task fMRI data or cFos stainings. BrainTrawler currently provides access to a small set of meta-analytic activation maps related to the terms “reward” and “pain” in the human brain, which were extracted from NeuroSynth, a platform for automated meta-analysis of neuroimaging data [13].

### Spatially resolved Transcriptomics Data

Transcriptomics data quantifies the so-called expression of genes in tissue. This information can be obtained through imaging methods, such as in-situ hybridization, or tissue sampling-based methods, such as single-cell RNA sequencing (scRNA seq.). In the former, expression is available for each voxel, while in the latter, expression is known for each sample location. This data is typically structured as a high-dimensional, sparse matrix where rows represent genes, columns represent samples, and each cell contains an expression value, accompanied by metadata such as sample tissue location, cell type, and subject information. In our application, we give users access to the extensive transcriptomics data resource of BrainTACO (see figure 2 by Ganglberger *et al*. [8]).

## WORKFLOWS AND REQUIREMENTS

We worked together with a team of domain experts from the pharmaceutical sector to identify tasks in the target gene identification workflow that our application should enable. These domain experts have thorough knowledge in drug target identification. The identified tasks can be broadly categorized into two categories: Circuit creation and DGEA based on circuits.

To investigate circuits that have already been described in the literature, neuroscientists should be able to recreate them in our application from circuit architecture descriptions (e.g. Beam *et al*. [12]). As an alternative, users should also be able to derive circuits directly from 3D activation signals (e.g., preprocessed fMRI activation maps or cFos stainings). In general, scientists should be able to flexibly adjust and customize the architecture of any circuit they have created.

Once the circuit has the desired properties, users should be able to use it for DGEA and see results quickly. Specifically, domain scientists should have the option to choose the entire circuit, but also specific nodes as query regions. Finally, neuroscientists should be able to filter resulting gene lists using reasonable statistical metrics and export them for downstream analyses, such as annotation of these genes using public databases (e.g., the OpenTargets Platform [14]).

We derived a list of requirements from the tasks described above. They can be summarized as follows:

- Users must be able to manually define circuits [R1];
- Users must be able to load circuits from a database [R2];
- Users must be able to generate circuits from brain activation signals (e.g., task fMRI) [R3];
- Users must be given anatomical context for circuits [R4];
- Users must be able to edit and save circuit properties [R5]
- Queries must use statistical testing similar to standard DGEA methods [R6];
- Queries must not take longer than few seconds to enable interactive exploration [R7];
- Queries must allow inspection on a node level [R8]
- Users must be able to interactively filter query results by statistical metrics [R9];

Requirements [R1-5] ensure that a flexible circuit concept is implemented, allowing multimodal circuit creation. The data mining approach for differentially expressed genes is constrained by [R6-8]. Finally, [R9] defines required interactions with generated DGEA results.

## FRAMEWORK OVERVIEW

Our framework can be broadly categorized into two main modules: Circuit Creation and Circuit Queries. A schematic depicting the interaction between these modules is shown in fig. 1.

**Figure 1:**
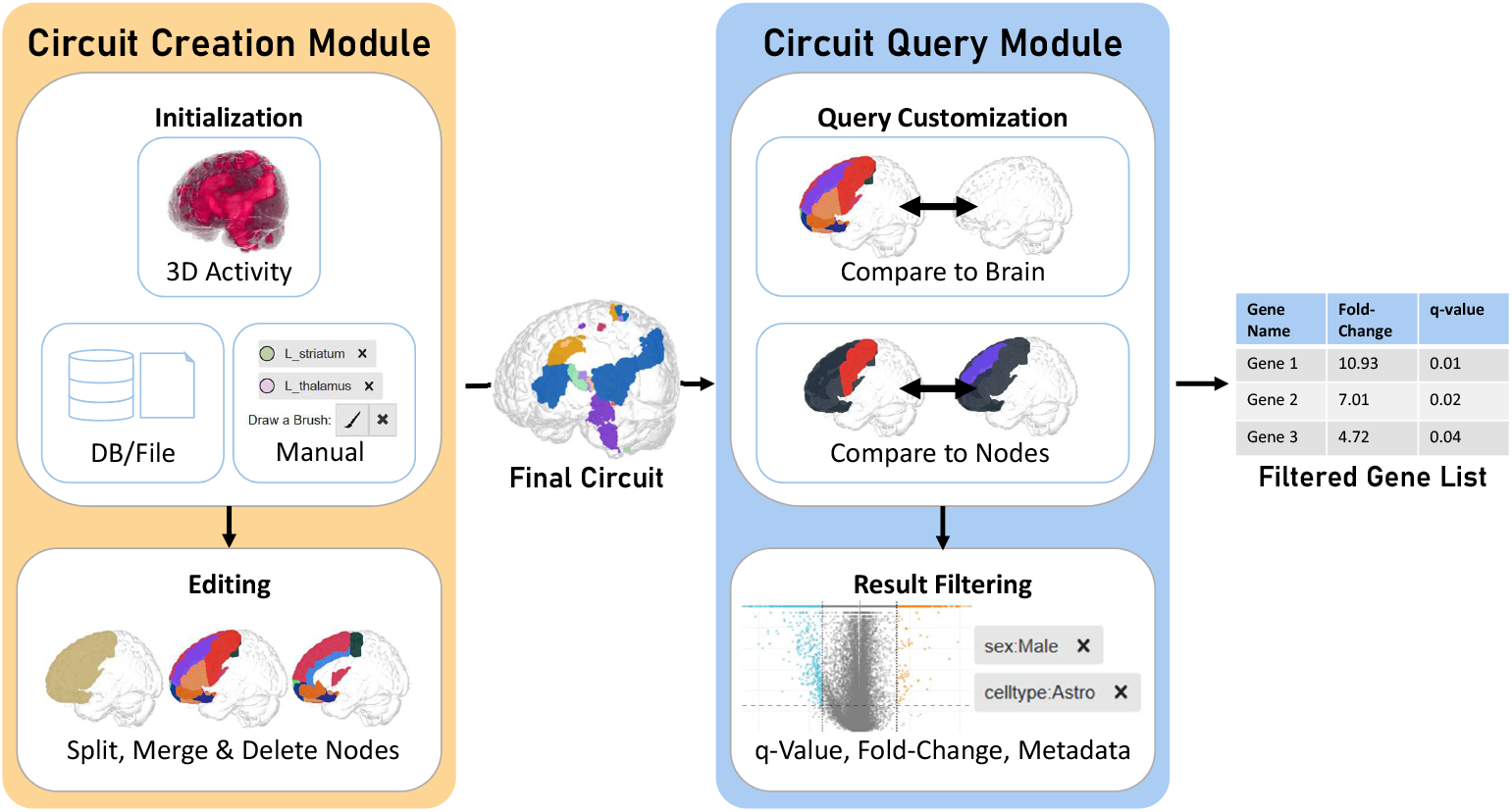
Schematic of the two modules in our framework. The circuit creation module (left) allows users to initialize and edit circuits. Circuits can then be used as the input for the circuit query module (right), where queries against the rest of the brain or specific circuit elements can be made. Result gene lists can be filtered by metadata as well as common metrics for specificity and statistical significance.

### Design process

The desired interactions and the design of our application were created in an iterative development and review cycle between developers and domain experts. We mainly used wireframe sketches for quickly demonstrating the user interface (UI) and planned interactions to users before writing code. During development, an intermediate state of the software was provided to domain experts for hands-on testing.

### Implementation and Availability

The presented approach was implemented as an extension of BrainTrawler [11], a web-based data exploration tool for multimodal brain data. BrainTrawler already provides access to a large collection of transcriptomics datasets for mouse and human [8]. BrainTrawler is a multi-component software, with a web-application written in Typescript/Javascript using React, a backend written in Go, and an OrientDB database. The spatial indexing framework was written in Go and operates as a standalone service.

A hosted and GDPR-compliant instance of the application including all mentioned public data is accessible to the public free of charge under https://braintrawlercircuit.vrvis.at/.

## Circuit Creation

The circuit creation module (see left part of fig. 1) allows users to initialize and edit arbitrary brain circuits. These circuits are later used as input for the circuit query module. An overview of the UI for circuit creation is shown in fig. 2a-d.

**Figure 2:**
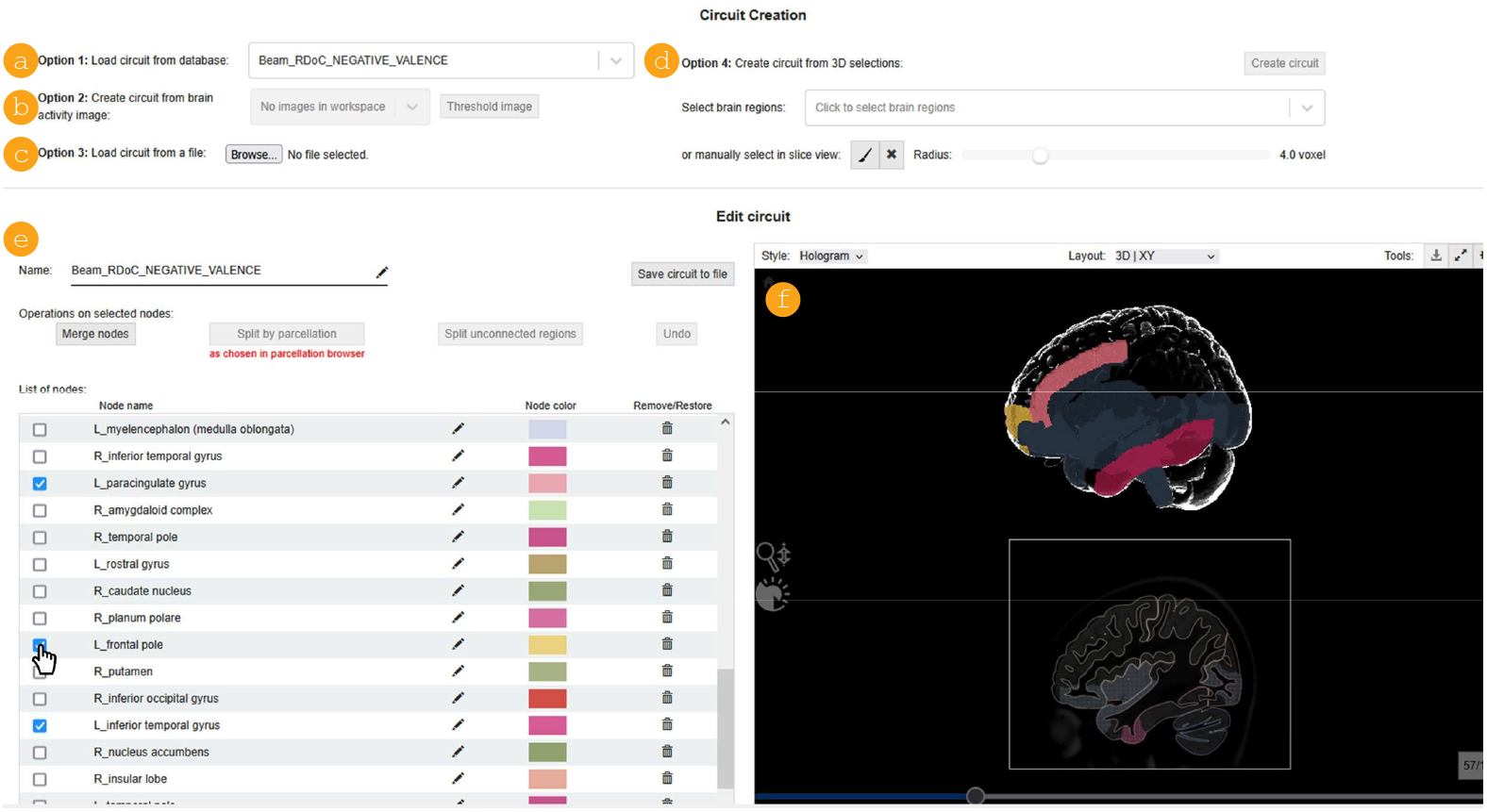
Circuit creation and edit UI. Circuits can be loaded from a database (a) or file (b), generated by applying a threshold to a brain activity signal (c) or manually defined (d). Once a circuit was created, it is visualized in the 3D and 2D renderer on the bottom right (f). The nodes “L_frontal pole”, “L_inferior temporal gyrus” and “L_paracingulate gyrus” are selected in the node list (e) and highlighted in the renderer.

### Circuit Initialization

We propose the following workflows for circuit initialization to meet the requirements: First, users can load predefined circuits from a database or from a file (see [R2] and fig. 2a&b). Descriptive metadata about each database circuit can be found in a separate page. Alternatively, activity-based circuits can be derived from any 3D brain activity signal (e.g., preprocessed task fMRI or cFos data, see [R3] and fig. 2c). To do this, the user performs image thresholding on the voxel-wise activity to generate a binary 3D image. This binary image is then divided into spatially disjoint volumes using a custom region growing algorithm. This algorithm visits the six nearest neighbors of a seed voxel, collects all active voxels in a group and then executes the same operation with the newly found active voxels. We consider all volumes smaller than 100 voxels for humans and 20 voxels for mice as not being relevant and discard them to reduce clutter. Then, each remaining volume is used as a node with a random color, as shown in fig. 3. Finally, users can manually define circuits by performing 3D brush selections and/or selecting brain regions from a standardized 3D brain parcellation (see [R1] and fig. 2d). These spatial selections are reflected in the 3D renderer and 2D slice view of the brain. The corresponding volumes are converted into a circuit by assigning each brain region to a node and collecting all brushes in a single node. Our reference atlases provide standard colors for each brain region, which are kept for the resulting nodes. Nodes from manual brushes are colored randomly.

**Figure 3:**
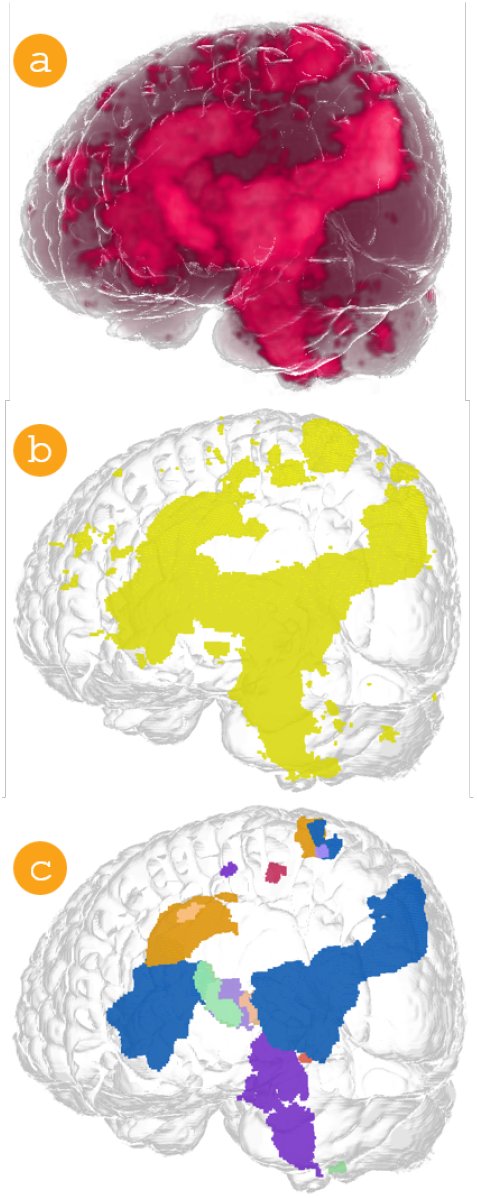
Example for creating a circuit from an activity signal (a). First, a threshold is applied to the 3D signal (b), then each spatially disjoint 3D volume above a fixed size threshold is considered as a node of the new circuit (c). Generated using BrainTrawler.

### Circuit Editing

After a circuit is initialized, it can be edited, investigated in 3D, and used for querying. The circuit edit UI is located directly below the circuit creation and shows a circuit overview with the list of nodes on the left and a 3D & 2D renderer visualizing the circuit on the right (see fig. 2e&f). In each list entry, we show a selection checkbox, an editable node name, the node color, and a button to delete the node. To facilitate detailed inspection of node locations beyond color-based identification, we use a focus-and-context-inspired approach. Initially, each node is displayed in a distinct color. When a node entry in the list is selected or hovered over, the corresponding node(s) remain in color within the renderer, while all others are grayed out. This hovering feature allows users to quickly pinpoint the location of a specific node. Clicking on a list entry selects a node, highlighting it permanently. Multiple nodes can be selected simultaneously, as illustrated in fig. 2e. The combination of hovering and selecting enables users to efficiently identify and group nodes for editing purposes.

Our framework provides four options for modifying nodes. Firstly, unwanted nodes can be deleted from the circuit by clicking the trash can icon in the node list (see bottom left in fig. 2). The deleted node is crossed out and removed from the renderer and queries. Secondly, nodes can be merged, with the resulting node containing the union of volumes of all input nodes. The merged node will inherit the names of all parent nodes. These two interactions are useful for reducing the complexity of a circuit. Since users were also interested in dissecting nodes to increase the spatial resolution of subsequent queries, we implemented the two interactions “split unconnected regions” and “split by parcellation”. With the first option, a node can be split into spatially disjoint volumes, each of which is assigned a new node. The same split operation is used in the process of generating a circuit from a brain activation signal, as shown in fig. 3. This operation is especially useful after a circuit was initialized through manual brushing, since all brushed voxels are initially collected in a single node. Brain parcellations are often organized hierarchically, i.e., brain regions can have sub-regions. Users can interact with these parcellations in our application and change the expansion state. Using “split by parcellation”, a node can be split along the boundaries of brain regions in the current expansion state (see fig. 4). This allows users to dynamically dissect a node into finer, anatomically motivated nodes for subsequent analysis steps. To convey anatomical context (see [R4]), the resulting nodes are named <ParentName>_<RegionName>, where RegionName is the brain region containing the node. In addition to editing the node architecture, users can modify the circuit name and node names of a circuit at any point. All edit operations are stored in the circuit history and can be reverted with the “Undo” button (see fig. 2). All circuits created with our framework can be exported to a file, preserving the circuit state as well as the history of edit operations for replicability (see [R5]). This option allows storing circuits between sessions and sharing them with colleagues.

**Figure 4:**
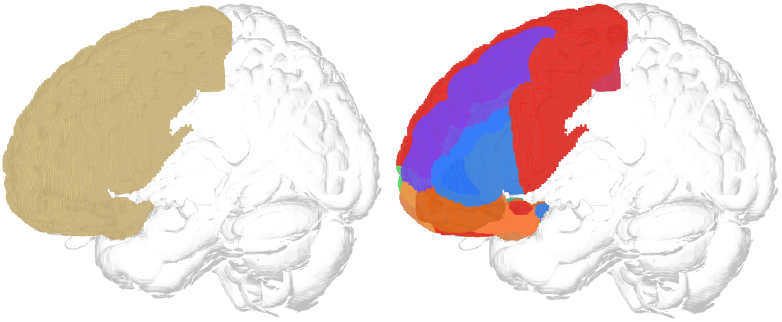
Example for splitting nodes along a brain parcellation. A node containing the frontal lobe is shown on the left, the result of “split by parcellation” is shown on the right. Generated using BrainTrawler.

## Circuit Queries

The second module of our application allows users to execute DGEA in the context of arbitrary brain circuits for various transcriptomics datasets. Input brain circuits are the result of workflows in the first module (see right part of fig. 1). The UI for the circuit query module is shown in fig. 5.

**Figure 5:**
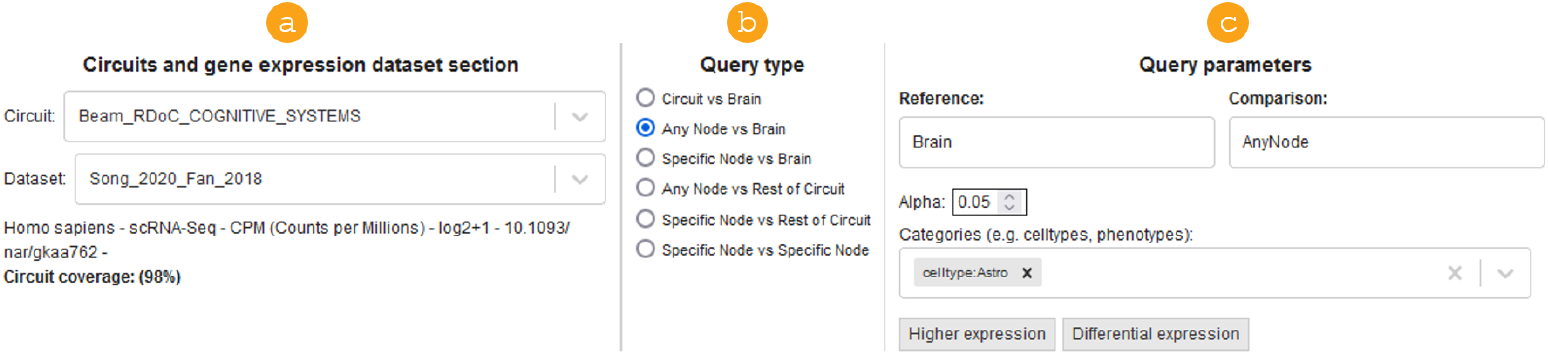
Circuit query user interface. After selecting a circuit and dataset (a), users can specify a query option Q1-Q6 (b). After the query type has been selected, the reference and comparison objects are shown on the right and query options, such as the threshold for statistical significance, and metadata filters can be set (c).

### Statistical Testing

The mathematical basis for circuit queries is a DGEA, which we implemented as Welch’s t-test combined with post-hoc corrections for multiple testing (see [R6]). We chose to implement t-tests because they are computationally inexpensive, which enables interactive queries (see [R7]). Results from this test are most reliable when the compared populations follow normal distributions. T-tests are commonly used for bulk RNA sequencing data, where normality is often assumed (see e.g., the Allen Human Brain Atlas API [15]). They have also been shown to perform similarly to more complex differential expression analysis methods in single-cell sequencing data [6], where the assumption of normality is usually violated.

To compute Welch’s t-test for a single gene between two query regions, the mean expression *μ*, variance of expression *σ*^2^, and the number of samples *n* in each region are required. To effectively extend our existing data storage and reading approach (see next section), we rewrite the variance as linear combinations of the sum of gene expression and the sum of squared gene expression. This allows calculating the variance without holding all expression values in the query region in memory. Let *x*_*i*_ be the gene expression of each sample/voxel *i* in the query area. Then, *μ* and *σ*^2^ are calculated according to

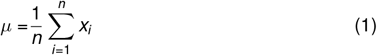

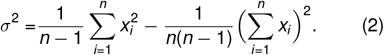

A derivation of this expression for *σ*^2^ is provided in the appendix. As a result of the t-test, the *p*-value for gene *i* can be obtained from the cumulative distribution function of a t-distribution with *ν* degrees of freedom at *x* = *t*_*test*_. Let *μ*_*j*_, 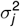, and *n*_*j*_ be mean, variance, and number of samples in the *j*^*th*^ query region (*j* = 1, 2).

Then

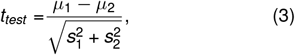

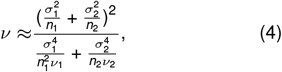

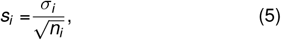

where *ν*_*i*_ = *n*_*i*_ *−* 1. We correct for multiple testing for the number of tested genes using the BenjaminiHochberg procedure for false discovery rate control. To do this, genes are ordered according to their *p*-value in ascending order. The corrected *p*-value (commonly referred to as q-value) for gene *i* is

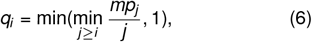

where *m* is the number of tested genes.

### Data Aggregation and Spatial Indexing

We use spatial indexing to efficiently retrieve *μ, σ*, and *n* for performing t-tests between arbitrary ROIs (see [R6]). To this end, we use the definitions shown in eq. (1) and eq. (2). Our work extends the approach taken by Ganglberger *et al*. [8], where only *μ* was encoded. We differentiate between sample-based gene expression data, such as RNA sequencing and microarray expression data, and volumetric data, such as in-situ hybridization expression data. In the first case, the expression of a gene is available only on the level of the entire tissue sample (e.g., a specific brain sub-region). In the latter case, a specific expression value is available per voxel for each measured gene. For obtaining the mean and variance of volumetric data in a query region, we use spatial indexing similar to Schulze [16]. For each voxel, the expression values *x* of all images at the voxel’s position are stored together (i.e., on the physical hard drive). Then, we order this per-voxel data along a space filling curve, which allows data in close 3D proximity to be stored in proximity on the hard-drive. This allows efficient, block-wise retrieval of *x* in the comparison/reference region, which is used to calculate *x* ^2^ and finally *μ* and *σ*for the t-test. The workflow for sample-based data is similar, but involves more pre-processing. Before reorganizing and indexing the data, we pre-aggregate samples with the same meta-data (e.g., dataset, cell type, sample location, subject age, etc.) so that 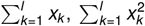 and *l* are known and stored for the aggregated samples, where *l* is the number of samples with similar meta-data. Pre-aggregation significantly improves query performance since it reduces the amount of samples that need to be queried at runtime by a factor of 10-100 (depending on the extent of the query) [8]. After this, we proceed, similarly to indexing imaging data, by ordering aggregated samples along a space-filling curve. This approach allows us to query the expression data needed for ttests in imaging and sample-based data for arbitrary ROIs in a few seconds.

### Query Workflow

Once the user is satisfied with a circuit, they use it as input for the circuit query module, as shown in fig. 1. The UI for this module is depicted in fig. 5. There, we call the first query region “comparison” and the second query region “reference”.

After choosing a transcriptomics dataset (see fig. 5a), the user can select a query type from six available options (Q1-Q6, see fig. 5a and tab. 1). These query types differ in the query regions used and allow the inspection of expression on a circuit and node level (see [R8]). The query options Q1-Q3 are intended for finding genes specific to the circuit or its nodes compared to the brain while options Q4-Q6 allow comparing expression between circuit nodes. To find genes that are specific to the entire circuit compared to the rest of the brain, users can execute a “Circuit vs. Brain” query (Q1). Similarly, genes that are specific to a single node can be found with a “Specific Node vs. Brain” query (Q3). For a more liberal search, users can choose an “Any Node vs. Brain” query (Q2), which lists all genes that are specific to any of the nodes in the circuit by executing a DGEA for each node. Similarly, the options “Any Node vs. Rest of Circuit” (Q4), “Specific Node vs. Rest of Circuit” (Q5) and “Specific Node vs. Specific Node” (Q6) can be used to inspect expression differences within a circuit. By combining these query options, users can obtain a detailed understanding of the transcriptional landscape of brain circuits.

**Table 1:** Explanation of the available query types for circuits. *B, C* and *c* denote the entire brain, the entire circuit and one of *n* circuit nodes, respectively.

|  | Query type | Usecase | Comparison | Reference |
| --- | --- | --- | --- | --- |
| Q1 | Circuit vs. Brain | Find genes specific to a circuit or circuit nodes. | $C$ | $B - C$ |
| Q2 | Any Node vs. Brain | | $c_i \forall i \in \{1, \dots, n\}$ | $B - C$ |
| Q3 | Specific Node vs. Brain | | $c_i, i \in \{1, \dots, n\}$ | $B - C$ |
| Q4 | Any Node vs. Rest of Circuit | Find genes specific to a node compared to other nodes. | $c_i \forall i \in \{1, \dots, n\}$ | $C - c_i$ |
| Q5 | Specific Node vs. Rest of Circuit | | $c_i, i \in \{1, \dots, n\}$ | $C - c_i$ |
| Q6 | Specific Node vs. Specific Node | | $c_i, i \in \{1, \dots, n\}$ | $c_j, j \in \{1, \dots, n\} \wedge j \neq i$ |

Once the dataset and query type is selected, the user can set a desired significance level (*α*) and select multiple metadata filters, e.g., specific cell types or subject ages to limit the query. To address the common issue of incomplete brain coverage in transcriptomics datasets, we provide a coverage summary in the query UI. This summary displays the percentage of volumetric overlap between the available expression data and the reference or comparison objects, allowing users to quickly assess data completeness for their regions of interest. Such information is especially important in query types Q1-Q3, where the rest of the brain (without the circuit) should serve as a reference object. Finally, users can execute queries either for higher expression (one-tailed tests) or differential expression (two-tailed tests). These queries use the implementation of DGEA described in the previous two sections. They return a list of genes, ranked by q-value and fold-change. For query options Q2 and Q4, one query is executed for each circuit node and the result lists are concatenated. We decided for this approach because domain experts pointed out that they expect a single result gene list that already includes node information rather than multiple lists. For each gene in the list, users can visualize 3D gene expression in the entire brain (see fig 6a), and inspect gene metadata and query results (see fig. 6c), such as the mean expression in the comparison and reference objects.

**Figure 6:**
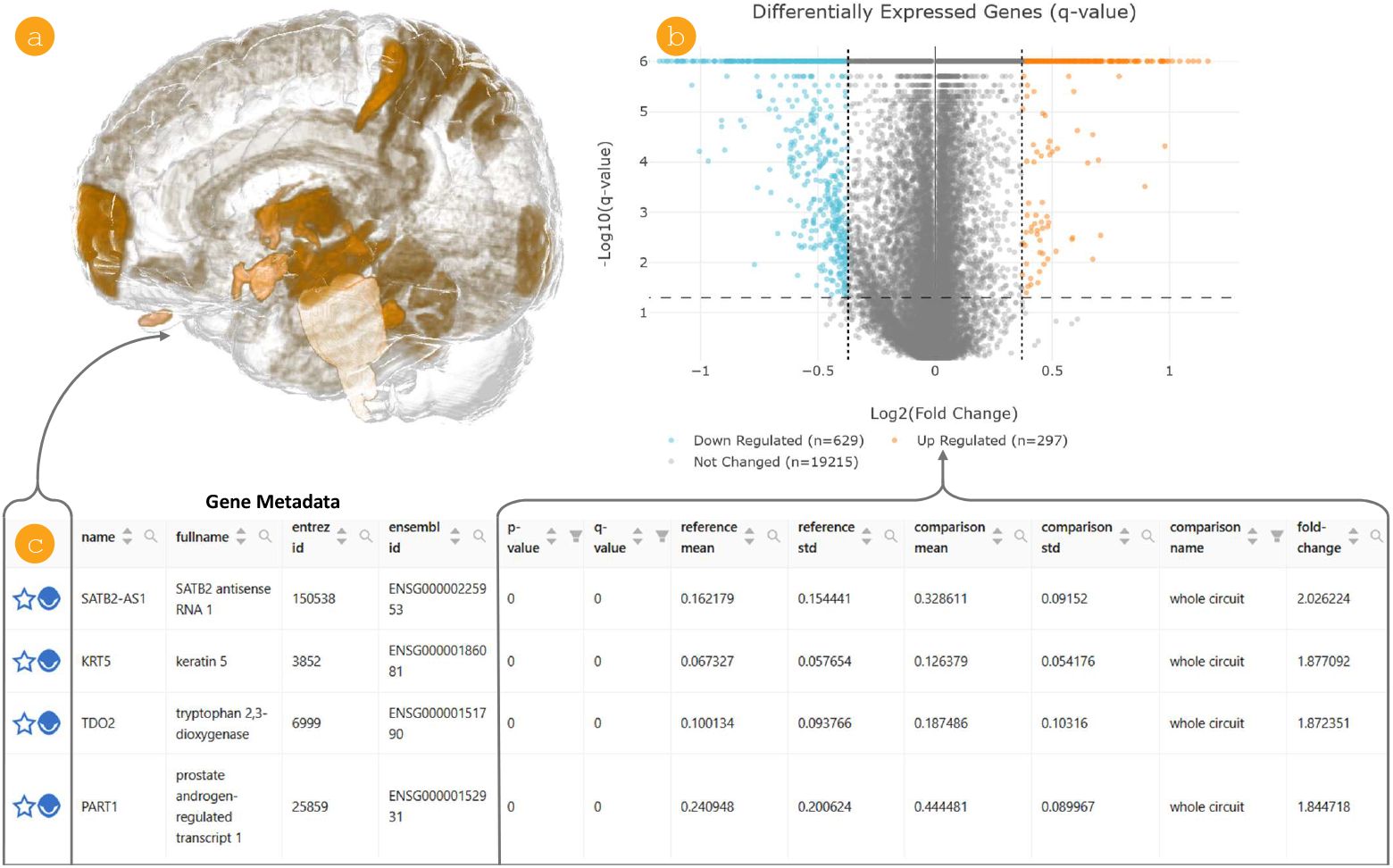
Example circuit query results. A result table (c) shows gene metadata and query results and allows users to visualize the expression of individual genes in 3D (a). The distribution of q-value, *p*-value and fold change is visualized using volcano plots (b). There, filters for *p*/q-value and fold change can be set interactively by dragging the dashed lines or entering values in an input field (not shown) to find specific and significantly expressed genes in the result table. Genes within the filter criteria are shown in blue/orange in the volcano plot.

### Result Filtering

*P*-value, q-value (*p*-value corrected for multiple testing), and fold-change are established metrics for statistical significance and gene expression specificity among domain experts. Our application allows users to filter result gene lists using these metrics (see [R9]). In addition to setting these values in input fields, they can also be set through interactive volcano plots that visualize the distribution of specificity metrics (see fig. 6). The volcano plot is a scatterplot with *x* = *log*_2_(fold-change) and *y* = *−log*_10_(q/p-value) and is commonly used to quickly identify entries that are both statistically significant and highly specific. This visualization is well-known in the user community, which reduces the need for visualization onboarding. In the application, users can set thresholds for the maximum *p*/q-value and minimum fold-change by specifying values in an input field or dragging the dashed threshold lines in the volcano plots. For the queries Q2 and Q4, which execute multiple DGEAs, we decided against using another visualization, such as scatterplot matrices, after discussions with domain experts. They pointed out that they perceive node information as secondary compared to the q-values and fold changes, which is why we did not highlight node differences in the visualization.

### Limitations

The querying approach presented here enables researchers without technical background to search for differentially expressed genes in arbitrary brain circuits. We chose to implement the DGEA as Welch’s t-test instead of more complex statistical texts, such as DESeq2 [5] or the Wilcoxon signed-rank test, to enable fast and interactive querying that is compatible with the existing spatial indexing framework (see [R7]). While this approach is computationally efficient, the underlying assumption of normality in the compared sample sets cannot be guaranteed, especially with scRNA seq. data. Nevertheless, t-tests are widely used for DGEA of microarray expression data, e.g., in the Allen Brain API [7], and were found to have similar reproducibility of top-ranked highly expressed genes in scRNA seq. data as methods better tailored towards scRNA seq. data [6]. Due to these limitations, we recommend using our software to build and refine hypotheses, followed by confirming these hypotheses with more rigorous statistical testing. Furthermore, not all transcriptomics datasets are useful for circuit queries, since many of them do not cover the whole brain and therefore often do not provide data in the comparison or reference regions. To address this issue, we provide a dataset coverage summary in the circuit query UI.

## USER FEEDBACK

To evaluate how well the requirements [R1-9] were met and as how useful users perceive the software, we conducted usability testing with five domain experts, loosely following a guide by Riihiaho [17]. We combined four established evaluation methods to obtain feedback from users: (i) observational task analysis, (ii) thinking aloud, (iii) qualitative feedback, and (iv) usability satisfaction evaluation (SUS [18]). Evaluation sessions lasted between 30–50 minutes. Two coauthors were present at every session, one acting as a moderator, while the second one took notes of user actions and responses. The evaluation process was structured as follows:

1. **Introduction:** We introduced the study purpose and procedure and explained the concept of thinking aloud. Users were asked if they agreed to the evaluation’s data processing, privacy, and anonymity terms. Then, we asked participants about their previous experience with BrainTrawler and the general process of target gene identification using DGEA. Afterwards, we explained the general concept of circuits and briefly presented the software interface.
2. **Task analysis:** We asked users to perform two tasks while thinking aloud. The moderator did not actively provide help, but gave hints in case the user was stuck. As a first task, we asked users to manually create a circuit, re-organize nodes, perform a query with this circuit and filter results by q-value and fold change. As a second task, we asked participants to load a specific brain activity image and create a circuit from it. The exact task instructions can be found in the appendix.
3. **Qualitative evaluation:** After finishing the tasks, we asked participants for general feedback on the application and task workflows and to provide suggestions for improvements. In particular, we asked them to compare these workflows to standard target gene identification processes and inquired if they trusted results obtained with our application.
4. **Usability survey:** Finally, we gathered subjective assessments of usability using an all-positive version of the SUS questionnaire [18] with a seven-point Likert scale. Participants were asked to fill the form after the video call to minimize the influence of the interview situation.

### Previous user experience

Four out of five participants reported having experience with previous versions of BrainTrawler before implementation of the extension involving circuits. One participant was not proficient in target gene identification and DGEA, but was aware of the basic principles. The domain experts reported that a standard DGEA workflow commonly involves multiple people and takes a minimum of two weeks to obtain a gene list for a given dataset, reference, and comparison object.

### Task analysis results

Three users quickly understood the manual circuit creation workflow while two others needed hints on the exact starting point (see fig. 2d). One user expected to be able to add additional nodes to a circuit once it was already created. The node list and merging operations were quickly understood by all users. While all users performed the correct query on their first try, one participant noted that they were not sure whether the chosen query type was indeed correct. All users managed to correctly filter the resulting gene list, but one participant was initially confused by the option of choosing the *α* value before executing the query, as instructions mentioned filtering for significant genes after the query. In the second task, two users were given a hint to find the option to create a circuit from a brain activity image (see fig. 2b).

### Qualitative feedback

All users with experience in DGEA approved that using t-tests is a valid choice, especially for exploration. Three participants were happy to have the possibility to quickly run differential queries without having to ask for help, stating that *‘the tool is a novel alternative to searching for existing literature on a circuit, because queries can be tailored to user needs’*. Those experts all mentioned that they plan to use our software to search for target genes in the coming months. Another user with a statistics background noted that they could imagine using the tool for high-level exploration before more rigorous statistical testing. The different ways of creating circuits were appreciated by multiple users, who stated that, for them, *‘circuit creation from neuroimaging data was not possible in the past’*. Four participants mentioned that the workflows and queries are organized reasonably, but several experts also noted that the amount of possible workflows can be confusing at the start. Participants noted that the node list and the linked 3D visualization were easy to interact with, and that results filtering using the volcano plot was useful. Two users reported initial difficulties in finding the manual circuit creation UI. One participant suggested adding abstract, results-focused workflow entry points, since the workflow goals of some tabs were not entirely clear to them. Another candidate reported feeling stuck after circuit creation and asked for more guidance on subsequent steps. Furthermore, multiple study participants mentioned that the application could be further improved by adding brain region information to the node list view. Finally, a participant mentioned that it would be great if circuits could also be used to query network data, such as functional or structural connectivity.

### System Usability Scale

All users filled out a System Usability Scale (SUS) questionnaire with a seven-point Likert scale (1-7). We calculated the SUS-score by subtracting 1 from each response, taking the sum, and then multiplying by 5*/*3 to bring the score to a range of 0-100 [18]. A summary of the results is shown in tab. 2. With the lowest individual SUS scores of approx. 70 and the highest result of approx. 98, all individual results lie within the adjective ratings “good” and “best imaginable” (see figure 13 in Bangor, Kortum and Miller [19]). Across all participants, the lowest ratings were given for S7 and S10, both of which refer to having to learn new concepts to use the application. This finding is consistent with statements given by some participants during the evaluation. However, since BrainTrawler is an expert software, a level of onboarding and additional necessary knowledge is expected. The best overall scores were obtained for statement S4 (*‘I think that I could use this system without the support of a technical person*.*’*).

After obtaining user feedback, we increased the visibility of the workflow entry points for circuit creation by marking and numbering each of them, as this seemed to cause confusion among users.

**Table 2:** Results of the SUS questionnaire. Result of individual questions can range from 1-7. Scores from 0-100 are calculated according to Sauro and Lewis [18]. “Good” scores exceed 70, while “excellent” scores exceed 85 [19]. Statements S1-S10 can be found in the appendix.

|  | S1 | S2 | S3 | S4 | S5 | S6 | S7 | S8 | S9 | S10 | SUS-score |
| --- | --- | --- | --- | --- | --- | --- | --- | --- | --- | --- | --- |
| P1 | 6.0 | 6.0 | 7.0 | 6.0 | 5.0 | 6.0 | 6.0 | 6.0 | 7.0 | 5.0 | 83.33 |
| P2 | 7.0 | 7.0 | 7.0 | 7.0 | 7.0 | 7.0 | 6.0 | 7.0 | 7.0 | 7.0 | 98.33 |
| P3 | 6.0 | 5.5 | 5.0 | 7.0 | 6.0 | 6.0 | 5.0 | 4.0 | 4.0 | 4.0 | 70.83 |
| P4 | 6.0 | 7.0 | 7.0 | 7.0 | 7.0 | 7.0 | 6.0 | 7.0 | 7.0 | 7.0 | 96.67 |
| P5 | 7.0 | 6.0 | 6.0 | 6.0 | 7.0 | 6.0 | 6.0 | 6.0 | 7.0 | 6.0 | 88.33 |

## CONCLUSION AND OUTLOOK

In this paper, we present a novel, workflow-driven approach for interactively mining transcriptomics data in brain circuits. We showcase a flexible approach to circuit creation that allows users to define circuits from multiple modalities. Furthermore, we present a performant solution for exploratory DGEA by customizing spatial indexing to t-tests, which allows differential queries between arbitrary ROIs at runtime. We evaluated the effectiveness of our approach through various types of user feedback and report high SUS scores and general satisfaction with the implemented workflows and concepts among all expert users. This software is the first to enable neuroscientists to perform DGEA using arbitrary brain circuits and nodes without the need for programming skills.

In future work, we will provide additional anatomical context by adding brain region information to the node list view, as suggested by domain experts. In addition, we will explore options for relating circuits to connectivity data, as suggested during the evaluation, e.g., by generating circuits from target or source queries or querying for the connectivity between nodes. In the current version of BrainTrawler, only a small set of human brain activation data is available. In the future, we plan to integrate extensive collections of such data, e.g., from NeuroSynth [13]. Enabling the direct upload of user-owned neuroimaging data, e.g., as NIfTI (.nii) files, would further enhance the usefulness of our tool. As mentioned in the limitations, many currently available datasets do not provide brainwide data. The integration of brain-wide expression atlases would allow users even more freedom in querying. It would be beneficial to add non-parametric tests, such as the Wilcoxon test, as an additional option for querying to increase statistical rigour, which would require an adapted data pre-processing and spatial indexing approach. Finally, the software could be further enhanced by annotating the resulting gene list with gene pathways by interfacing with public pathway databases.

While our application focuses on querying transcriptomics data in the brain, it might also be useful in other fields that rely on DGEA, such as cancer research. In general, the querying module can be extended beyond transcriptomics to work with multiple spatial *omics* data, such as ATAC sequencing or proteomics. The concept of circuits is quite specific to the field of neuroscience, but our approach of combining spatial indexing with statistical testing could be used in any field that uses statistics on large image collections, such as materials science.

## Supporting information

User study tasks

Derivation of Variance

System usability scale

## ACKNOWLEDGMENTS

We thank Theo Widmer and Alina Jaud for their contribution to the interactive volcano plots. The VRVis GmbH is funded by BMK, BMAW, Tyrol, Vorarlberg, and Vienna Business Agency in the scope of COMET - Competence Centers for Excellent Technologies (911654) which is managed by FFG.

**Tobias Peherstorfer** is a researcher in the Biomedical Image Informatics research group at VRVis GmbH. His current research interests are brain data integration and multimodal brain data analyses. He received his Master’s degree in Physics from TU Wien. He is the corresponding author of this article. Contact him at.

**Sophia Ulonska** is a senior researcher in the Biomedical Image Informatics research group at VRVis GmbH, Austria. She received the PhD in the doctoral program Technical Science, field of studies Chemical and Process Engineering, at TU Wien.

**Bianca Burger** is a senior researcher in the Biomedical Image Informatics research group at VRVis GmbH. She received the Ph.D. degree from the Medical University of Vienna in the doctoral program Medical Imaging with her work focusing on topics in Computational Neuroscience.

**Simone Lucato** is a researcher in the Biomedical Image Informatics research group at VRVis GmbH, Austria. He received his B.Sc. in Computer Science at University of Padova.

**Bader Al-Hamdan** is a researcher in the Biomedical Image Informatics research group at VRVis GmbH, Austria. He received his B.Sc. in Computer Science at University of Jordan.

**Marvin Kleinlehner** is a researcher in the Biomedical Image Informatics research group at VRVis GmbH, Austria. He is currently pursuing a B.Sc. in Media Informatics and Visual Computing at TU Wien.

**Till F. M. Andlauer** is a Principal Scientist in Boehringer Ingelheim’s Computational Innovation department. He received his doctor’s degree in Biomedicine at the University of Würzburg and completed his Habilitation in Molecular Medicine at the School of Medicine of the Technical University of Munich.

**Katja Bühler** is scientific director of VRVis GmbH, Austria, and the group lead of the Biomedical Image Informatics research group. She received her doctor’s degree in Computer Science at TU Wien. Contact her at. Additional Declarations: The authors declare no competing interests.

