## Supplementary material for "Circuit Mining in Transcriptomics Data": User study tasks

### BrainTrawler User study tasks

Before starting the user study, please open the following link:
https://braintrawler-circuit.vrvis.at/

Log in with the following credentials:

User name:
reviewer
Password:
t4K69|5wjI4)j;0>LVJn3eXQ

#### **Task 1:**

##### Topic: Creating circuits manually and querying

Please create a circuit in the human brain using the brain regions "Thalamus", "Striatum" and "cingulate gyrus, rostral (anterior) part" for both brain hemispheres.

Then, group mirrored brain regions in the same node (one node should contain left and right Thalamus).

Use this circuit to search the Allen Human Brain Atlas (Hawrylycz_2012) for genes that are highly expressed in the entire circuit.

How many genes are highly expressed in this circuit with a q-value $\leq$ 0.03 and a fold-change $\geq$ 4?

**Task 2:**

#### Topic: Creating circuits from activity signals

Please reload the application by pressing F5 or clicking the reload icon in your browser.

Then, please add a 3D brain activity image related to the term "painful" to your workspace from the "Browse Database" tab.

Then, create an activity-based circuit from it in the "Circuit Creation" page with a threshold value of 4.

How many nodes are created?
