## Supplementary material for "Circuit Mining in Transcriptomics Data": Derivation of Variance

### Derivation of variance representation

In our publication, we use the following identities for mean $\mu$ and variance $\sigma^2$:

$$\mu = \frac{1}{n}\sum_{i=1}^{n} x_i \tag{1}$$

$$\sigma^2 = \frac{1}{n-1}\sum_{i=1}^{n} x_i^2 - \frac{1}{n(n-1)}(\sum_{i=1}^{n} x_i)^2. \tag{2}$$

To see, that $\sigma 2$ can be written in this way, consider the following rearrangement of the usual formula:

$$\sigma^2 = \frac{1}{n-1}\sum_{i=1}^{n}(x_i - \mu)^2 = \frac{1}{n-1}\sum_{i=1}^{n}(x_i^2 - 2x_i\mu + \mu^2) \tag{3}$$

$$= \frac{1}{n-1}(\sum_{i=1}^{n} x_i^2 - 2\mu\sum_{i=1}^{n} x_i \cdot \frac{n}{n} + n\mu^2) \tag{4}$$

$$= \frac{1}{n-1}(\sum_{i=1}^{n} x_i^2 - 2n\mu^2 + n\mu^2) = \frac{1}{n-1}(\sum_{i=1}^{n} x_i^2 - n\mu^2) \tag{5}$$

$$= \frac{1}{n-1}\sum_{i=1}^{n} x_i^2 - \frac{n}{n-1}\frac{1}{n^2}(\sum_{i=1}^{n} x_i)^2 = \frac{1}{n-1}\sum_{i=1}^{n} x_i^2 - \frac{1}{n(n-1)}(\sum_{i=1}^{n} x_i)^2 \tag{6}$$
