## Supplementary material for "Circuit Mining in Transcriptomics Data": System usability scale

|  | Strongly agree |  |  | Neutral |  |  | Strongly disagree |
| --- | --- | --- | --- | --- | --- | --- | --- |
| I think that I would like to use this system frequently | ☐ | ☐ | ☐ | ☐ | ☐ | ☐ | ☐ |
| I found the system to be simple | ☐ | ☐ | ☐ | ☐ | ☐ | ☐ | ☐ |
| I thought the system was easy to use | ☐ | ☐ | ☐ | ☐ | ☐ | ☐ | ☐ |
| I think that I could use this system without the support of a technical person | ☐ | ☐ | ☐ | ☐ | ☐ | ☐ | ☐ |
| I found the various functions in this system were well integrated | ☐ | ☐ | ☐ | ☐ | ☐ | ☐ | ☐ |
| I thought there was a lot of consistency in this system | ☐ | ☐ | ☐ | ☐ | ☐ | ☐ | ☐ |
| I would imagine that most people would learn to use this system very quickly | ☐ | ☐ | ☐ | ☐ | ☐ | ☐ | ☐ |
| I found the system very intuitive | ☐ | ☐ | ☐ | ☐ | ☐ | ☐ | ☐ |
| I felt very confident using the system | ☐ | ☐ | ☐ | ☐ | ☐ | ☐ | ☐ |
| I could use this system without having to learn anything new | ☐ | ☐ | ☐ | ☐ | ☐ | ☐ | ☐ |
